# Rbm10 facilitates heterochromatin assembly via the Clr6 HDAC complex

**DOI:** 10.1101/518936

**Authors:** Martina Weigt, Qingsong Gao, Hyoju Ban, Haijin He, Guido Mastrobuoni, Stefan Kempa, Wei Chen, Fei Li

**Affiliations:** Laboratory for Functional Genomics and Systems Biology, Berlin Institute for Medical Systems Biology, Max-Delbrück-Center for Molecular Medicine, Berlin 13125, Germany; Department of Biology, New York University, New York, NY 10003-6688, USA; Integrative Metabolomics and Proteomics, Berlin Institute of Medical Systems Biology, Max-Delbrueck Center for Molecular Medicine, Berlin 13125, Germany; Department of Biology, Southern University of Science and Technology, Shenzhen, Guangdong, China; Medi-X Institute, SUSTech Academy for Advanced Interdisciplinary Studies, Southern University of Science and Technology, Shenzhen, Guangdong, China

**Keywords:** epigenetics, splicing factor, *Schizosaccharomyces pombe*, histone deacetylase, H3K9 methylation

## Abstract

Splicing factors have recently been shown to be involved in heterochromatin formation, but their role in controlling heterochromatin structure and function remains poorly understood. In this study, we identified a fission yeast homologue of human splicing factor RBM10, which has been linked to TARP syndrome. Overexpression of Rbm10 in fission yeast leads to strong global intron retention. Rbm10 also interacts with splicing factors in a pattern resembling that of human RBM10, suggesting that the function of Rbm10 as a splicing regulator is conserved. Surprisingly, our deep-sequencing data showed that deletion of Rbm10 caused only minor effect on genome-wide gene expression and splicing. However, the mutant displays severe heterochromatin defects. Further analyses indicated that the heterochromatin defects in the mutant did not result from mis-splicing of heterochromatin factors. Our proteomic data revealed that Rbm10 associates with the histone deacetylase Clr6 complex and chromatin remodelers known to be important for heterochromatin silencing. Deletion of Rbm10 results in significant reduction of Clr6 in heterochromatin. Our work together with previous findings further suggests that different splicing subunits may play distinct roles in heterochromatin regulation.

## Introduction

In eukaryotic cells, DNA and histones are organized into the highly-ordered chromatin structure. Chromatin forms two distinct domains, euchromatin and heterochromatin. Euchromatin is less condensed and associated with actively transcribed genes, whereas heterochromatin is tightly coiled and transcriptionally inactive. Heterochromatin usually consists of highly repetitive sequences, which are commonly found around pericentromeres and telomeres. Heterochromatin plays essential roles in gene expression, chromosome segregation and genome stability (1–3).

The pattern of histone modifications in heterochromatin contributes to the organization of the chromatin domain. Heterochromatin is characterized by hypoacetylation of histone lysines and methylation of histone H3 on lysine 9 (H3K9me) (1,3). Histone deacetylases (HDACs) are responsible for hypoacetylation in heterochromatin, while H3K9me is catalyzed by the histone methyltransferase SUV39, an enzyme conserved from yeasts to humans. H3K9me serves as a binding site for heterochromatin protein 1 (HP1), which recruits additional factors to mediate heterochromatin assembly (1,3). Small interfering RNAs (siRNAs) generated by RNA interference pathway have been shown to be important for heterochromatin structure and function in a variety of organisms (1).

The fission yeast *Schizosaccharomyces pombe* has emerged as an excellent model organism for understanding heterochromatin formation. It harbors three constitutive heterochromatic regions: the pericentromere, telomere and mating type region. These regions are enriched with H3K9me, which is recognized by the human HP1 homolog, Swi6. H3K9me in fission yeast heterochromatin is regulated by the multi-subunit CLRC complex, which contains Clr4, the H3K9 methyltransferase, Rik1, Dos1/Raf1, Dos2/Raf2, Cul4 and Lid2, a H3K4 demethylase (4–9). H3K9 methylation in heterochromatin also requires RNAi.

Heterochromatin is transcribed during S phase of the cell cycle (10,11). The heterochromatin transcripts are subsequently processed into siRNAs by Ago1, the RDRC complex and Dicer. The siRNAs are loaded into RITS complex (Ago1, Chp1 and Tas3), which in turn guides the CLRC complex to heterochromatin regions (1,12,13) to mediate H3K9me. The DNA Pol epsilon subunit, Cdc20, also contributes to the recruitment of CLRC complex during the S phase (14–16).

Like other eukaryotes, histones in fission yeast heterochromatin are hypoacetylated. Fission yeast contains all three subtypes of histone deacetylases (HDACs): Class I, Class II and Class III. Clr6, a Class I HDAC, is a homolog of mammalian HDAC1 and HDAC2, and deacetylate multiple lysine residues on histone H3 and H4. Clr3 is a Class II HDAC, responsible for H3K14 deacetylation (17–19). Sir2 is a conserved member of the Sirtuin family of Class III HDACs that uses NAD^+^ as a cofactor (20,21). All of these HDAC contributes to heterochromatin silencing in fission yeast (17,19). Our recent results reveal that the histone fold subunit of the DNA Pol epsilon, Dpb4, is important for recruitment of Sir2 during DNA replication (22).

Recently, splicing factors have been linked to heterochromatin formation in fission yeast. But the mechanisms for how these splicing factors participate in heterochromatin formation remains unclear. A prevailing model proposes that splicing factors, independently from splicing, directly interacts with RNAi components to mediate the recruitments of the CLRC complex to heterochromatic regions (23). Another study suggests that transcripts derived from pericentromeric repeats contain introns, and splicing factors may mediate the splicing of the intron of the heterochromatin transcripts, which in turn influences siRNA generation (24). A more recent study by Kallegren, et al. suggested that splicing factors may contribute to heterochromatin assembly indirectly by regulating the proper splicing of heterochromatin factors (25).

RBM10 has been recently characterized as a new member of splicing factors, and associated with lung cancer and congenital disorders, such as the TARP syndrome and XLMR (26–29). RBM10 contains five distinct domains, including two RNA recognition motifs (RRM), two zinc fingers and one G patch motif. Splicing involves an ordered, stepwise rearrangement of protein and RNA content in splicesome. An early step in spliceosome assembly is the formation of the prespliceosome (complex A). The prespliceosome rearranges to the precatalytic spliceosome (complex B), which subsequently gives rise to the activated spliceosome (complex B*). Complex B* carries out the first catalytic step of splicing, generating complex C, which in turn carries out the second catalytic step (30). It has been shown that RBM10 is associated with prespliceosome (complex A) and activated spliceosome (complex B*) in human cells (31–35). The human RBM10 has been shown to regulate exon skipping by repressing the splicing of introns (26). In addition to its role in splicing, human RBM10 was also found to be part of histone H2A deubiquitinase complex (36), suggesting a role in linking splicing and chromatin regulation.

In this study, we identified a fission yeast homologue of RBM10, which was named *rbm10*^+^. Overexpression of Rbm10 in fission yeast results in strong growth defects and global intron retention. We further demonstrated that Rbm10 interacts with heterochromatin factors, including Clr6 complex and chromatin remodelers, suggesting a previously unrecognized mechanism underlying splicing factor-mediated heterochromatin silencing.

## Results

### Human splicing factor RBM10 is conserved in *S. pombe*

Via homolog search, we identified SPAC57A7.13, an uncharacterized protein, as the homolog of human RBM10 in *S. pombe*. We hereafter named it Rbm10. The human RBM10 and its *S. pombe* homolog share an amino acid identity of 26% (Supplementary Figure S1). Despite relatively short length of Rbm10 in fission yeast, all five domains are well conserved, namely, two zinc-finger domains, two RRM-motifs and a G-patch domain (Figure 1A). Majority of Zinc-finger domains functions as interaction modules that bind DNA, RNA or proteins (37,38), whereas RRM motifs and the G-patch domain usually associate with RNA (39). To determine the cellular distribution of Rbm10, we constructed the GFP-tagged *rbm10*^+^ under its native promoter and replaced the endogenous *rbm10*^+^. Cells carrying Rbm10-GFP has no detectable defect, indicating that Rbm10-GFP is functional. We found that, similar to human RBM10, Rbm10-GFP is enriched in the nucleus (Figure 1B).

**Figure 1.**
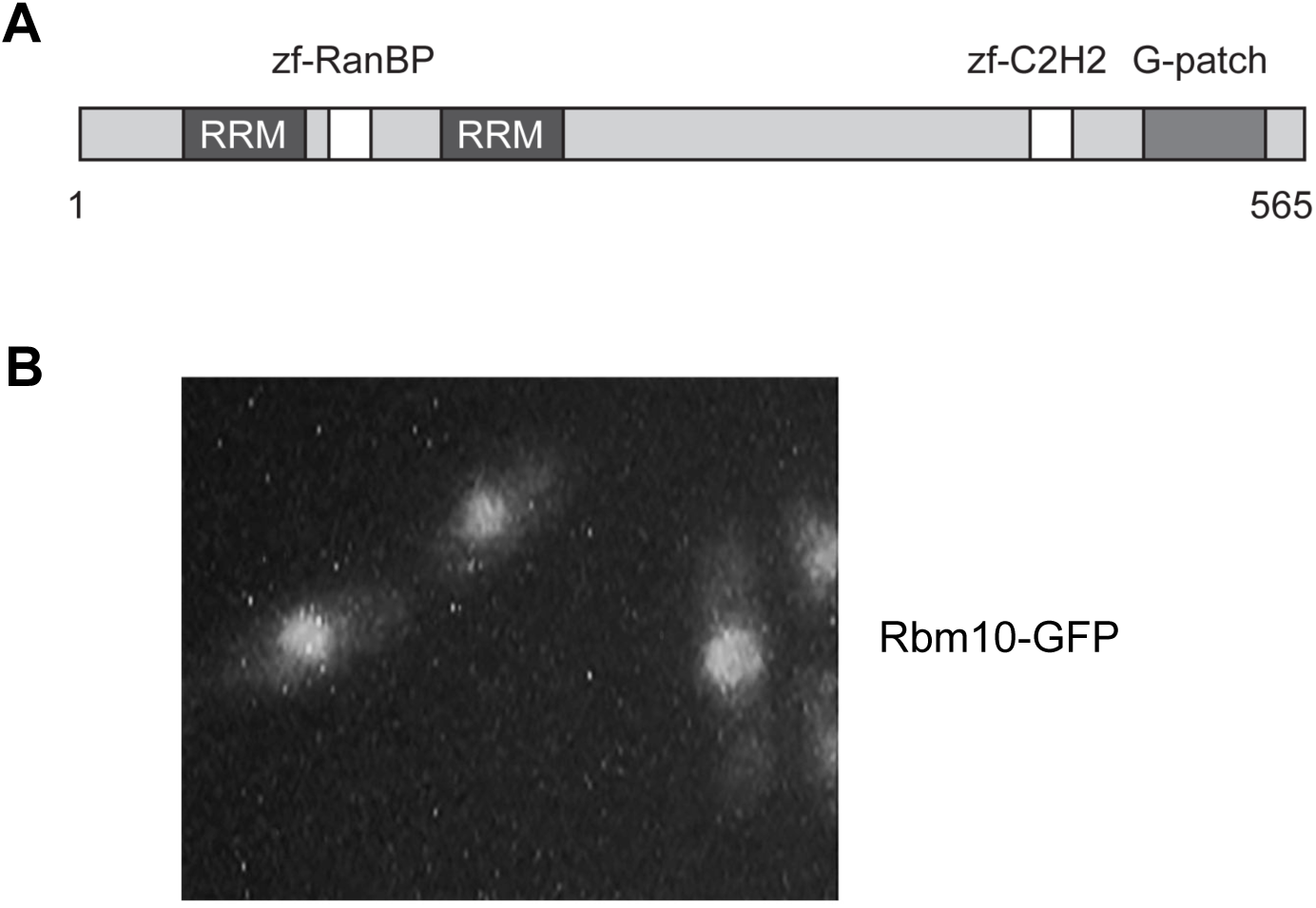
Human RBM10 is conserved in *S. pombe*. (**A**) Schematic diagram showing the domain structure of Rbm10 in *S. pombe*. (**B**) Rbm10 is enriched in the nucleus. Rbm10-GFP was constructed at its native locus under the control of its own promoter.

### Overexpression of Rbm10 leads to massive intron retention

To characterize the function of Rbm10 in fission yeast, we constructed *rbm10*^+^ under the strong, inducible *nmt1* promoter and overexpress it in wild-type cells. The *nmt1* promoter is induced in the absence of thiamine. We found that overexpression of Rbm10 in wild type results in slow growth (Figure 2A). In addition to the strong growth defects, cells overexpressing Rbm10 display aberrant cell shapes. They are often massively elongated or branched in to Y-like shapes. The nuclei in these cells also appear to be abnormal. Multiple fragmented nuclei were frequently observed (Figure 2B). Given that *rbm10*Δ showed only a mild phenotype as decribed below, we were surprised to find the observed toxicity resulting from overexpression of Rbm10. To gain insight into the molecular nature of Rbm10 overexpression, we performed unbiased genome-wide RNA-seq using cells overexpressing Rbm10. Cells overexpressing for 20 hours were used to minimize secondary effects. We detected massive changes in intron retention from the RNA-seq data: 2263 introns showed increased retention (Figure 2C and Supplementary Table S1), whereas 6 introns have decreased retention (Figure 2C and Supplementary Table S2). These data demonstrated that Rbm10’s function as a splicing regulator is conserved in fission yeast, which can be observed at least under overexpression condition.

**Figure 2.**
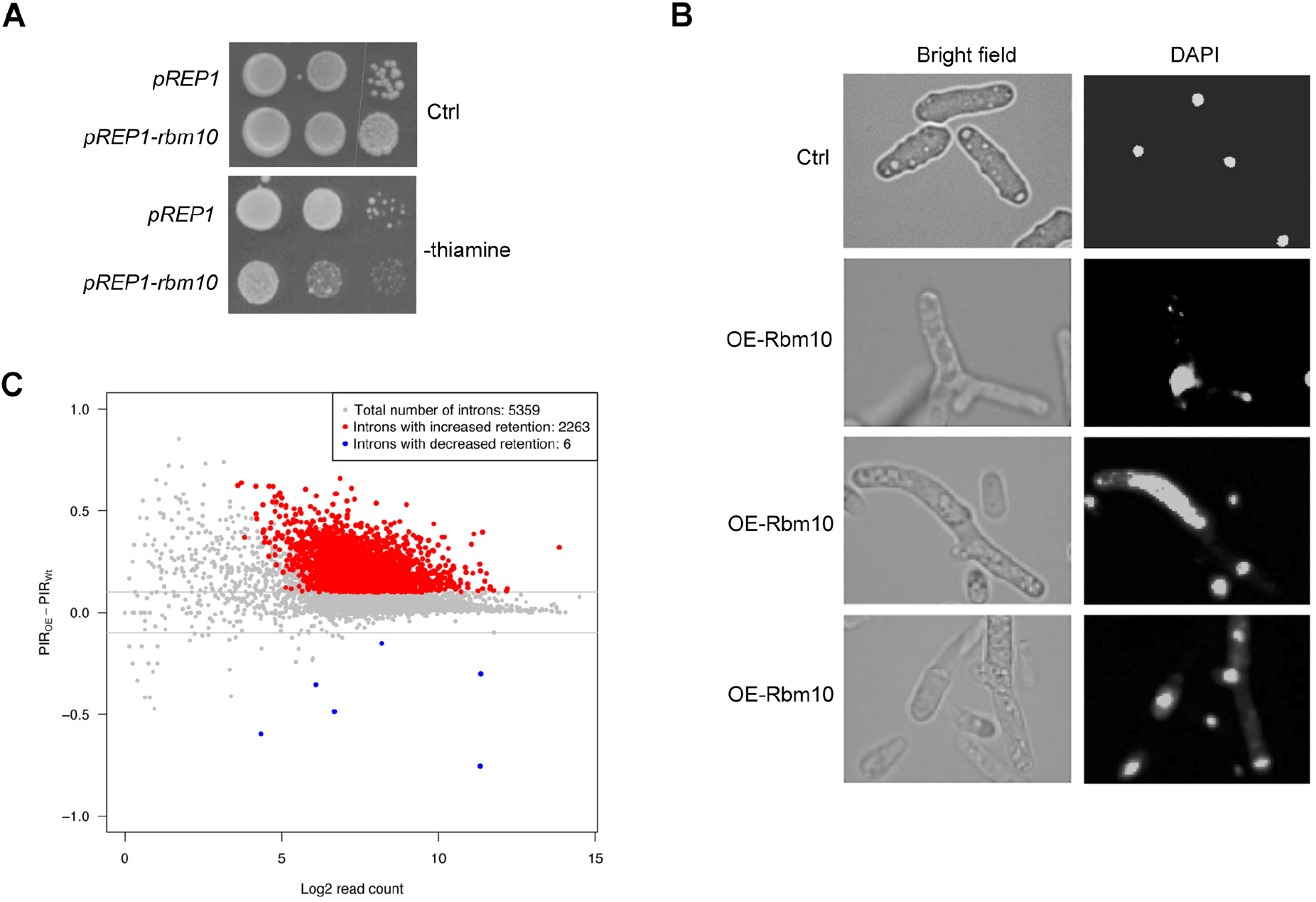
Overexpression of Rbm10 results in severe growth defects and massive intron retention. **(A)** Cells overexpressing Rbm10 display slow growth. Serial dilutions of indicated cells carrying pREP1-*rbm10* were plated in the PMG minimal medium without thiamine. Empty pREP1 vector was used as a control. Ctrl, a control plate containing thiamine. (**B**) Cells overexpressing Rbm10 showed abnormal cell shapes and nuclear organization. Left, bright-field microscopic images. Right, DAPI staining. OE, overexpression. (**C**) MA plot comparing the intron retention patterns between *rbm10*-overexpressing cells and WT. Log2 transformed read count for each event was plotted on the X axis, and the PIR difference was on the Y axis. The red and blue dots represented the events with significantly different intron retention patterns. The grey lines indicate 10% PIR difference.

### Deletion of *rbm10*^+^ induces minor changes in gene expression and intron retention

To further understand the role of Rbm10, we constructed a deletion strain, *rbm10*Δ, in which the complete gene was replaced with a kanamycin selection cassette via homologues recombination. *rbm10*Δ cells are viable and do not show obvious growth defects, indicating that Rbm10 is not essential for viability.

To investigate how Rbm10 affects gene expression and splicing, we performed genome-wide RNA-seq in the *rbm10*Δ mutant. We reproducibly found that out of 6748 genes, the expressions of 292 genes in *rbm10*Δ cells were increased compared to wild type (Figure 3A and Supplementary Table S3), whereas the gene expressions of 99 genes were reduced in the mutant (Figure 3A and Supplementary Table S4). However, the fold-changes of these gene expressions were very modest, with only 21 (0.3%) genes whose expression altered more than 2 fold. Our RNA-seq also allowed us to analyze the effect of *rbm10*Δ mutant on the intron splicing. Out of 5357 expressed introns, we detected 38 intron retention events in *rbm10*Δ mutant (Percent Intron Retained difference (ΔPIR) > 10%, Figure 3B and Supplementary Table S5). We also found that 7 intron splicing have been enhanced (ΔPIR < 10%, Figure 3B and Supplementary Table S6). Again, the majority of these changes were affected to a modest level. Previous studies have shown that alternative splicing exists in fission yeast, although not very common (40). Given that human RBM10 is involved in alternative exon usage (41), we also examined alternative exon usage in *rbm10*Δ mutant. However, our results show that no case with significant difference between WT and *rbm10*Δ mutant can be detected (Figure 3C). Taken together, our results show that deletion of Rbm10 has minor effect on gene expression and splicing.

**Figure 3.**
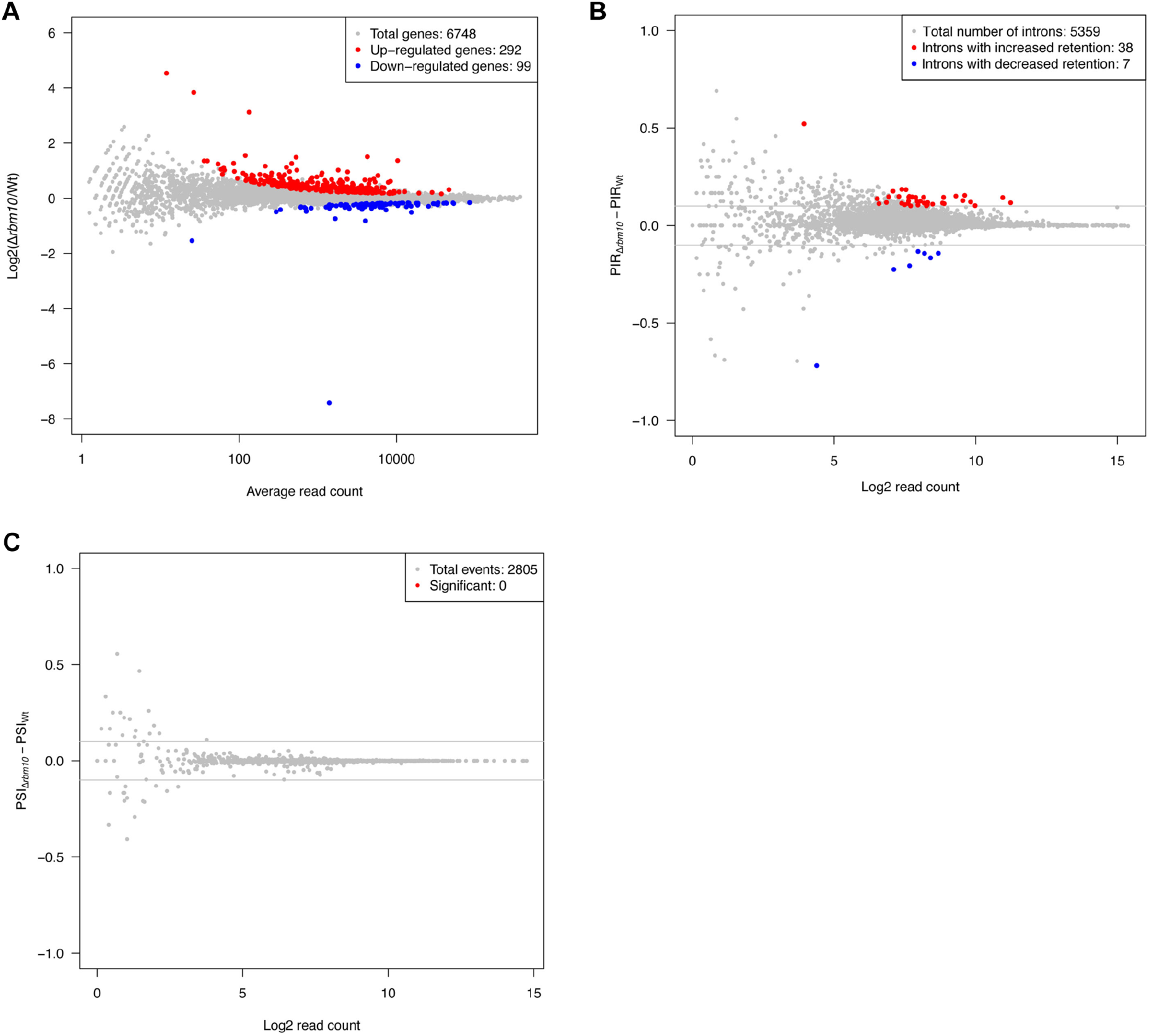
Deletion of Rbm10 has minor effect on gene expression and intron retention. (**A**) MA plot comparing the gene expression patterns between *rbm10*Δ cells and WT. The average read count for each gene was plotted on the X axis, and the log2 transformed fold change was on the Y axis. The red and blue dots represented the genes with significantly different gene expression patterns. (**B**) MA plot comparing the intron retention patterns between *rbm10*Δ cells and WT. Log2 transformed read count for each event was plotted on the X axis, and the PIR difference was on the Y axis. The red and blue dots represented the events with significantly different intron retention patterns. The grey lines indicate 10% PIR difference. (**C**) MA plot comparing the exon skipping patterns between *rbm10*Δ cells and WT. Log2 transformed read count for each event was plotted on the X axis, and the Percent Spliced In difference (ΔPSI) was on the Y axis. The red dots represented the events with significantly different exon skipping patterns. The grey lines indicate 10% PIR (Percent Intron Retained) difference.

Wang et al. proposed that human RBM10 promotes exon skipping via binding in the vicinity of splice sides, resulting in delaying their splicing choice; thus, these exons with relatively weaker splice side become more excluded during alternative splicing (26). However, alternative splicing is a very rare event in fission yeast (40). It is likely that a delay in splicing resulting from Rbm10 splicing activity does not manifest in alternative splicing, but rather in intron retention. Our data showing a dramatic increase in intron retention in *S. pombe* cells overexpressing Rbm10 is consistent with this idea.

### Deletion of *rbm10*^+^ leads to impaired heterochromatin silencing

Splicing factors have been implicated in heterochromatin assembly. We next characterized how Rbm10 affects heterochromatin status. Fission yeast harbors three constitutive heterochromatic regions, namely pericentromeres, telomeres and the mating type locus. To analyze pericentromeric silencing in the *rbm10*Δ mutant, a wild type (WT) strain carrying a *ura4*^+^ reporter integrated in the pericentromeric outer repeat (*otr*) was crossed with the mutant. The resulting *rbm10*Δ cells with *ura4*^+^ reporter at *otr* region were analyzed by growth assays in the medium without uracil. The *rbm10*Δ cells grow significantly faster than WT (Figure 4A), indicating that the pericentromeric silencing in the mutant is disrupted. The strains were further analyzed in the media containing 5-fluoro-orotic acid (FOA), which is toxic to cells expressing uracil. WT cells exhibit strong silencing in *otr* region, as indicated by their robust growth in the FOA media. The slow growth of the *rbm10*Δ cells in the FOA media (Figure 4A) confirmed that the silencing at pericentromeric heterochromatin in *rbm10*Δ mutant is lost.

**Figure 4.**
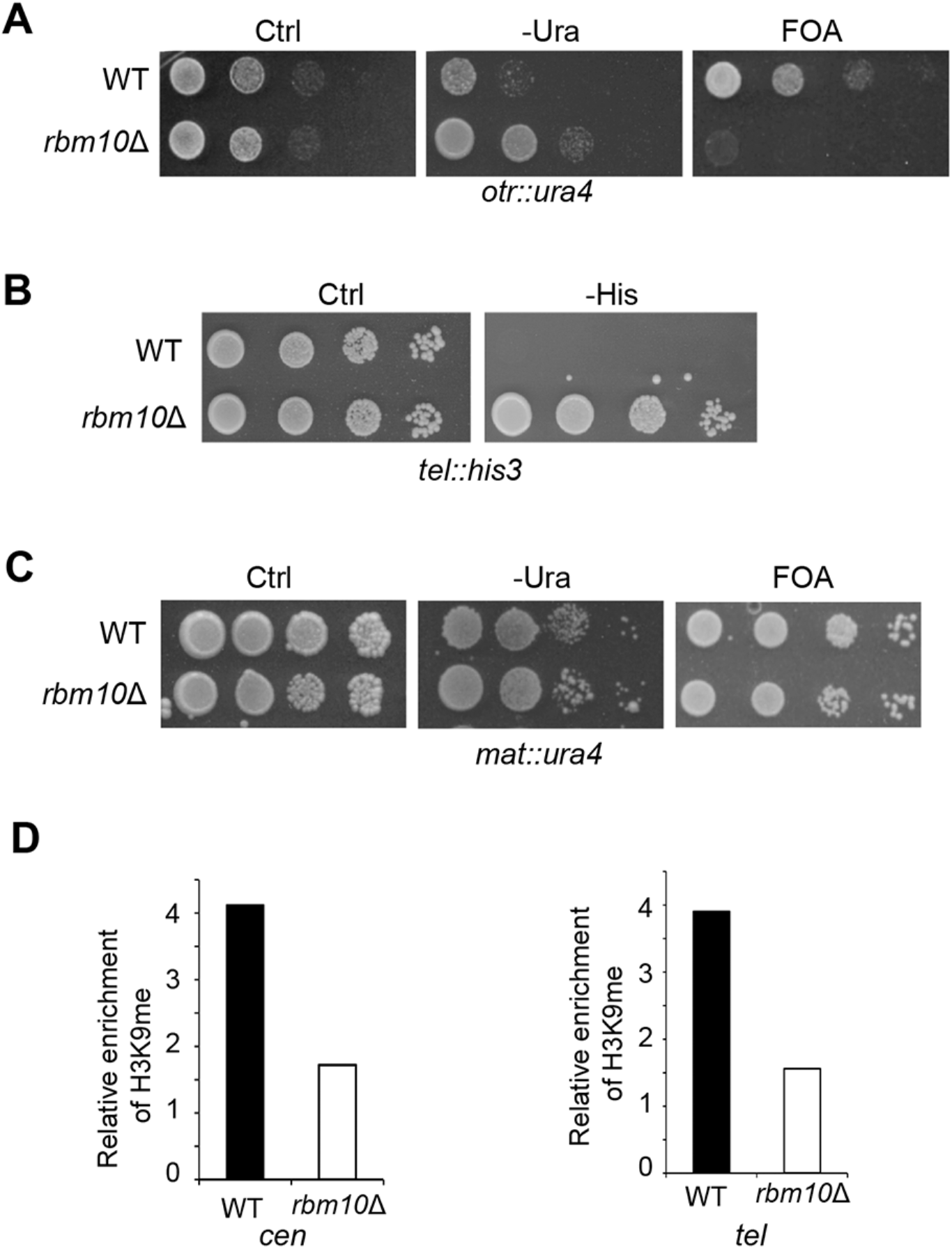
Rbm10 is important for heterochromatin silencing. (**A**) Serial dilutions of indicated cells harboring *ura4*^+^ at the *otr* region were plated in the PMG minimal medium without uracil (-Ura) or the EMM medium with the counter-selective FOA. Ctrl, a control plate containing uracil. WT, wild type. (**B**) Serial dilutions of *rbm10*Δ cells and WT as control carrying *his3*^+^ at the telomeric region (*tel*) were plated in the PMG medium without histidine (-His). (**C**) Growth assays of *rbm10*Δ and WT cells carrying *ura4*^+^ at the mating-type region in the PMG medium without uracil (-Ura) or with the FOA media. (**D**) Analysis of H3K9 methylation in the *otr* region by ChIP. ChIP assays were performed using an antibody against H3K9 di-methylation and primers specific for in the *otr* region. *act1*^+^ was used as a control.

We also examined the heterochromatin silencing in telomere and mating type locus in *rbm10*Δ using *his3*^+^ and *ura4*^+^ reporters inserted to these regions, respectively. Our growth assays using minimum media without histidine demonstrated that silencing in telomere is strongly lost in *rbm10*Δ cells (Figure 4B). However, *rbm10*Δ cells with *ura4*^+^ in mating type locus did not show major growth deficiency relative to WT in minimum media without uracil and the counter-selective FOA medium (Figure 4C), indicating that the heterochromatin silencing in this region is not significantly affected by deletion of *rbm10*^+^. This different effect between centromere and mating type heterochromatin in *rbm10*Δ is likely due to the redundant pathways involved in heterochromatin formation in mating-type locus (42,43).

We next analyzed H3K9me in centromeres and telomeres using ChIP. We observed that H3K9me level is reduced to approximately 50% of WT at both peri-centromeres and telomeres in *rbm10*Δ (Figure 4D). The loss of H3K9me at peri-centromeres and telomeres is consistent with our growth assays, indicating that heterochromatin silencing is impaired in these regions in *rbm10*Δ.

It has been shown that mutants of splicing factors, such as Cwf14, can induce mis-splicing of mRNA transcripts from heterochromatin factors, including Ago1, which in turn results in heterochromatin defect (25). However, our transcriptome profiling from the *rbm10*Δ mutant showed the minor differences in *rbm10*Δ in intron retention, indicating that a secondary effect by *rbm10*Δ via the splicing of heterochromatin factors is unlikely to be the primary cause for heterochromatin defects.

### Rbm10 interacts with Clr6 and is important for recruitment of Clr6 to heterochromatin

To gain more insight into the function of Rbm10, we determined to identify its interacting proteins using the Tandem Affinity Purification (TAP) method. For this, we constructed a TAP tag with a combination of FLAG and HA tags at the N-terminus of *rbm10*^+^ at its endogenous locus. FLAG-HA-Rbm10 is fully functional, as it does not cause any silencing defects in the pericentromeric *otr* region (Figure 5A). However, we observed that the expression level of endogenous FLAG-HA-Rbm10 is extremely low (Data not shown). We thus constructed FLAG-HA-Rbm10 under the *nmt1* promoter, and overexpressed FLAG-HA-Rbm10 for TAP tag purification. We collected cells within 24 hours of induction in order to minimize the potential secondary effects. We performed a two-step affinity purification of FLAG-HA-Rbm10 and subjected the sample to mass spectrometry (MS) analysis (Figure 5B). From three independent affinity purifications, we obtained 853 interacting proteins (Figure 5C and Supplementary Table S7) found in all replicates but being absent from control purifications. These proteins included factors involved in heterochromatin silencing, chromatin remodeling, and RNA processing, and transcription (Figure 5D and 6A). GO Term analysis with PANTHER further revealed an overrepresentation of RNA metabolic process (p=9.75×10^−5^), chromatin organization (p=2.2×10^−16^) and DNA-dependent transcription (p=1.06×10^−2^).

**Figure 5.**
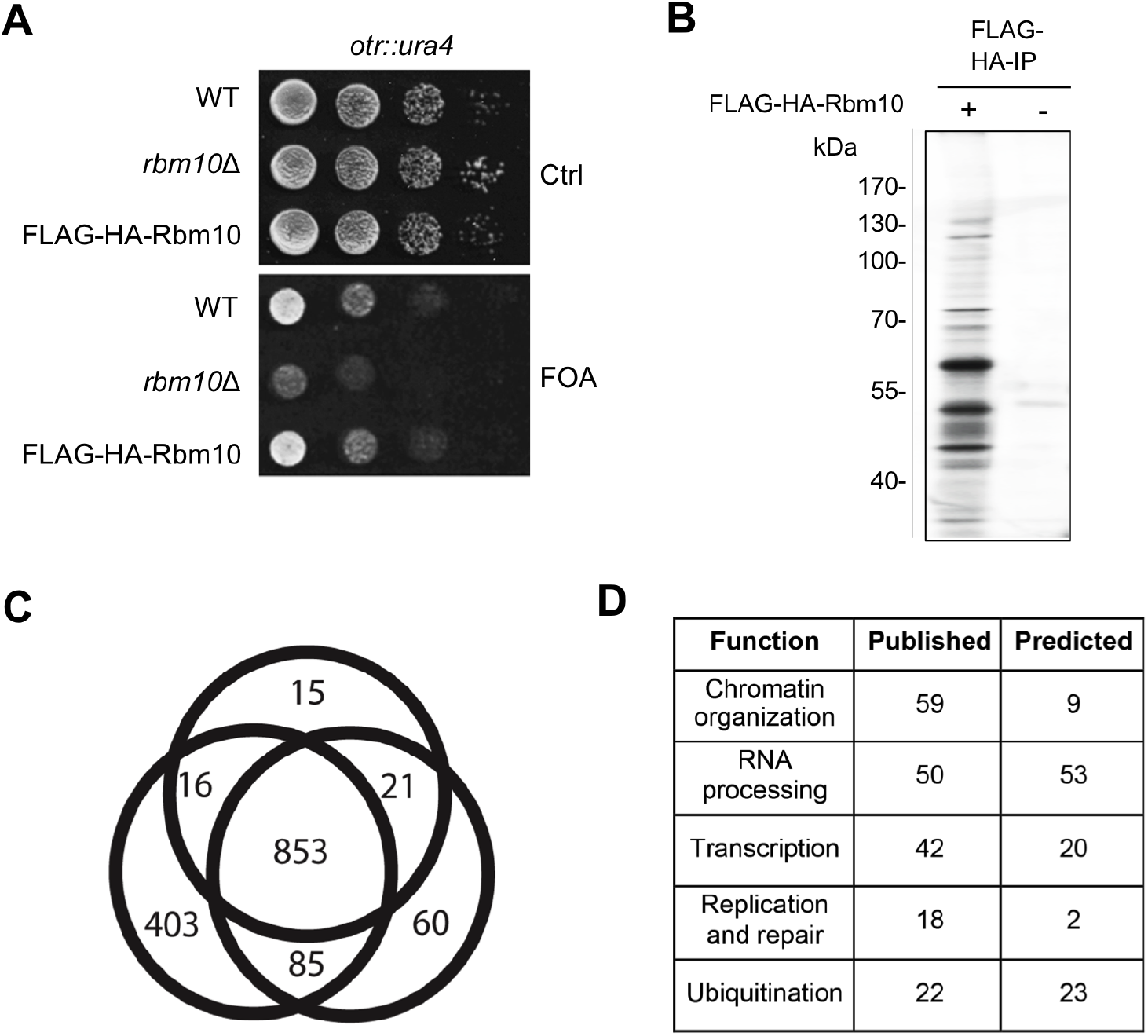
Mass spectrometry analysis of TAP-tag purified Rbm10. (**A**) Cells carrying FLAG-HA tagged Rbm10 under its own promoter expressed from its endogenous locus display no detectable silencing defect. Serial dilutions of indicated cells harboring *ura4*^+^ at the *otr* region were plated in the EMM medium containing FOA. Ctrl, a control plate without FOA. (**B**) A silver-stained gel showing the TAP-tag purification of Rbm10 and a control purification from an untagged strain. (**C**) Venn diagram showing the overlap of proteins detected by mass spectrometry. (**D**) The proteins identified by mass spectrometry were grouped based on their functions, as indicated in the table.

**Figure 6.**
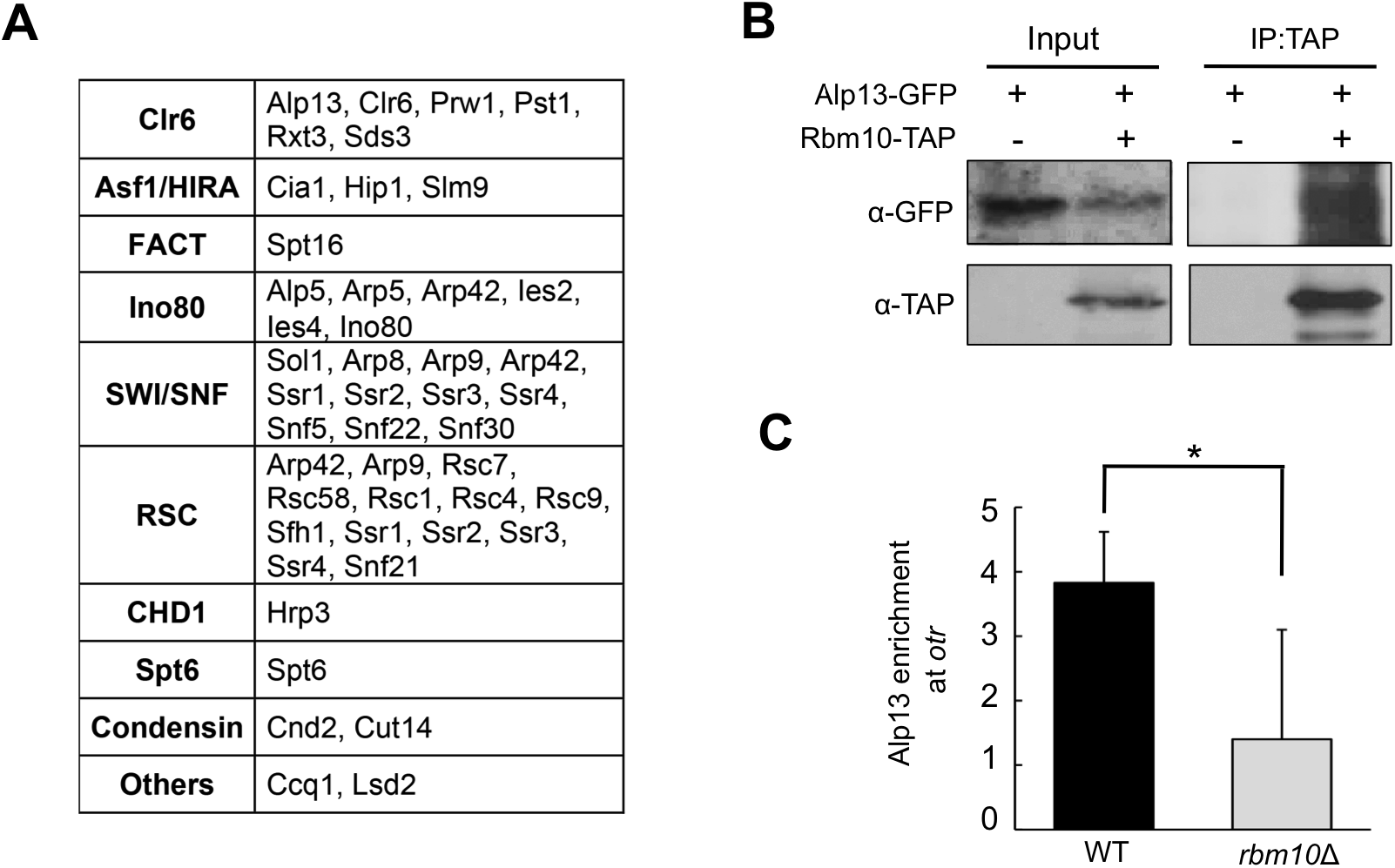
Rbm10 interacts with Clr6 and chromatin remodeling complexes. (**A**) Components of HDAC Clr6 and chromatin remodeling complexes identified by mass spectrometry analysis of Rbm10 were listed. Condensin components and other known heterochromatin factors identified were also shown. (**B**) Cell lysates from cells expressing Alp13-GFP and Rbm10-TAP under their own promoter at their native sites were immunoprecipitated with an antibody specific for TAP, and analyzed by immunoblotting using a GFP antibody. Cells expressing Alp13-GFP alone were used as a control. (**C**) Analysis of the association of Alp13-GFP in the *otr* region in the indicated strains by ChIP. ChIP assays were performed using an antibody against GFP. *act1*^+^ was used as a control. Error bars indicate SD. (*) P < 0.05

Clr6 HDAC complexes are important for heterochromatin silencing, and fall in two main classes I and II in fission yeast (18). Five members from the HDAC Clr6 I complex and also Alp13 from the Clr6 II complex can be found to be associated with Rbm10 (Figure 6A and Supplementary Table S7). Particularly, Prw1, a component of the Clr6 complex I, achieves the highest percentage of sequence coverage identified from our MS data.

We note that all the TAP purification experiments were performed using cells expressing FLAG-HA-Rbm10 to a higher level than endogenous Rbm10. To rule out the possibility that the interactions we identified are the artifacts caused by overexpression of Rbm10, we performed co-immunoprecipitation (Co-IP) to validate the interactions of Rbm10-TAP with Alp13-GFP, both of whom are expressed under their own promoter at the endogenous sites.

It has been shown that Alp13 is important for heterochromatin silencing (44). Rbm10-TAP and Alp13-GFP do not causes any detectable defect (data not shown) and are thus functional. Our co-IP experiments confirmed the interaction of Rbm10 with Alp13 (Figure 6B). To determine how Rbm10 affects the recruitment of the Clr6 complex to heterochromatin, we analyzed the *rbm10*Δ mutant carrying Alp13-GFP by ChIP assays. Our results showed that Alp13-GFP in the pericentromeric *otr* region is significantly reduced in *rbm10*Δ, indicating that Rbm10 is important for recruitment of the Clr6 complex to heterochromatin (Figure 6C).

In addition, we found that Rbm10 is copurified with Asf1/Cia1 and the HIRA histone chaperone complex (Figure 6A). It has been shown that Asf1 interacts with HIRA to mediate heterochromatin silencing (45,46). We also found that Rbm10 associates with FACT, Spt6 and CHD1/Hrp3 chromatin remodelers. These chromatin remodeling factors have also been shown to be important for heterochromatin formation (47–49). We detected a strong link of Rbm10 with RSC, SWI/SNF, and Ino80 chromatin remodeling complexes. All thirteen members of the RSC complex and eleven of twelve members of the SWI/SNF complex were found in the MS results (Figure 6A and Supplementary Table S7). We also found that condensin components in the Rbm10-purified product. Condensin has been implicated in pericentromeric heterochromatin assembly in fission yeast (10,50,51). In addition, Rbm10 interacts with Uhp1, an ubiquitinated histone-like protein, and Lsd2, a conserved histone demethylase, both of which are linked to heterochromatin formation (52,53). Rbm10 is also copurified together with Ccq1, which is required for silencing in telomere and facultative heterochromatin (54,55). Interestingly, Rbm10 associates with transcription factors, including all five members of the PAF complex, and the transcriptional activator Byr3, which was detected with a sequence coverage of 65% and the highest intensity to mass ratio in the whole dataset. Heterochromatin in fission yeast can be transcribed during S phase of the cell cycle, and the non-coding transcripts are processed into siRNAs by RNAi machinery (10,14). These transcription factors may be involved in regulation of heterochromatin transcription.

### Rbm10 interacts with splicing factors

Our MS analysis also detected more than 30 splicing factors interacting with Rbm10 (Table 1). Most of them are from the spliceosome A and B components, such as U1 snRNP, Sf3a and Sf3b, U2AF and NTC complex. This finding is in concordance with interaction data from human RBM10, which has been shown to be part of the spliceosome A and B complex (31–33). These results demonstrate that Rbm10 interaction with the spliceosome are conserved between human and fission yeast. Our data further indicate that, even though *rbm10*Δ mutant shows only modest changes in splicing, Rbm10 in fission yeast nevertheless is involved in RNA splicing, consistent with the Rbm10 overexpression data (Figure 3).

**Table 1.** Splicing factors that Rbm10 interacts with

| ID | Name | Description |
| --- | --- | --- |
| SPAC4D7.13 | Usp104 | U1 snRNP-associated protein Usp104 |
| SPBC4B4.09 | Usp105 | U1 snRNP-associated protein Usp105 |
| SPBC839.10 | Usp107 | U1 snRNP-associated protein Usp107 |
| SPAP8A3.06 | Uaf2 | U2AF small subunit, U2AF-23 |
| SPAC9.03c | Brr2 | U5 snRNP complex subunit Brr2 |
| SPAC644.12 | Cdc5 | cell division control protein, splicing factor Cdc5 |
| SPBC19C2.01 | Cdc28 | ATP-dependent RNA helicase Prp8 |
| SPBC16A3.18 | Cip1 | RNA-binding protein Cip1 |
| SPBC215.12 | Cwf10 | U5 snRNP GTPase subunit Cwf10 |
| SPCP1E11.07c | Cwf18 | complexed with Cdc5 protein Cwf18 |
| SPAC3A12.11c | Cwf2 | RNA-binding protein Cwf2 |
| SPBC211.02c | Cwf3 | complexed with Cdc5 protein Cwf3 |
| SPBC31F10.11c | Cwf4 | complexed with Cdc5 protein Cwf4 |
| SPCC550.02c | Cwf5 | RNA-binding protein Cwf5 |
| SPBC354.12 | Gpd3 | glyceraldehyde 3-phosphate dehydrogenase Gpd3 |
| SPBC29A10.10c | Dbl8 | tRNA-splicing endonuclease positive effector Dbl8 |
| SPAC23H3.02c | Ini1 | RING finger-like protein Ini1 |
| SPBC6B1.07 | Prp1 | U4/U6 x U5 tri-snRNP complex subunit Prp1 |
| SPAC27F1.09c | Prp10 | U2 snRNP-associated protein Sap155 |
| SPCC10H11.01 | Prp11 | ATP-dependent RNA helicase Prp11 |
| SPAPJ698.03c | Prp12 | U2 snRNP-associated protein Sap130 |
| SPAC29A4.08c | Prp19 | ubiquitin-protein ligase E4 |
| SPBC1861.04c | Prp24 | RNA-binding protein Prp24 (predicted) |
| SPBC119.13c | Prp31 | U4/U6 x U5 tri-snRNP complex subunit Prp31 |
| SPBC16H5.10c | Prp43 | ATP-dependent RNA helicase Prp43 |
| SPBP22H7.07 | Prp5 | WD repeat protein Prp5 |
| SPAC227.12 | Rna4 | U4/U6 x U5 tri-snRNP complex subunit Prp4 family, Rna4 |
| SPAC19G12.07c | Rsd1 | RNA-binding protein Rsd1 (predicted) |
| SPAC22A12.09c | Sap114 | U2 snRNP subunit Sap114 |
| SPBC29A3.07c | Sap14 | U2 snRNP-associated protein SF3B14 Sap14 |
| SPAC22F8.10c | Sap145 | U2 snRNP-associated protein Sap145 |
| SPBC36.09 | Sap61 | U2 snRNP-associated protein sap61 |
| SPAC4F8.12c | Spp42 | U5 snRNP complex subunit Spp42 |
| SPAC4D7.13 | Usp104 | U1 snRNP-associated protein Usp104 |
| SPBC4B4.09 | Usp105 | U1 snRNP-associated protein Usp105 |
| SPBC839.10 | Usp107 | U1 snRNP-associated protein Usp107 |

## Discussion

Previous studies have shown that human RBM10 is an RNA splicing protein, responsible for the X-linked TARP syndrome, although the mechanisms by which Rbm10 causes the disease remains unclear (26,27,29). In this study, we identified its homologue Rbm10 in fission yeast. We showed that Rbm10 in fission yeast interacts with a variety of splicing factors, and overexpression of Rbm10 results in strong intron retention. These data suggest that the role of Rbm10 in splicing regulation is conserved. Surprisingly, our genome-wide RNA analysis showed that deletion of *rbm10*^+^ in fission yeast results in minor splicing defects. However, heterochromatin formation is severely disrupted in *rbm10* mutant. Our data strongly support that abnormality in heterochromatin formation in the mutant arises independently from defects in splicing. We further demonstrated that Rbm10 interacts with heterochromatin factors, including Clr6 complex and chromatin remodelers, suggesting a novel mechanism underlying splicing factor-mediated heterochromatin silencing. Our findings reveal previously unrecognized features of Rbm10 in chromatin regulation, and may help elucidate its roles in human disease.

### Role of Rbm10 in heterochromatin formation

As in mammals, heterochromatin in fission yeast is enriched with hypoacetylation of histones and H3K9 methylation. These heterochromatin hallmarks are instrumental for heterochromatin structure and function (1,3). In fission yeast, siRNAs generated from heterochromatin transcripts during S phase promotes H3K9me by guiding the CLRC complex containing the H3K9 methyltransferase, Clr4, to heterochromatin (10,11,14,56). It has been shown that splicing factors are required for RNAi-mediated heterochromatin assembly in fission yeast. However, the mechanism behind the role of splicing factors in heterochromatin assembly remains poorly understood. There is an ongoing discussion in the field as to whether splicing factors participate in heterochromatin formation directly. A study by Bayne *et al*. has shown that in splicing factor mutants, such as *prp10-1*, splicing and heterochromatin defects can be uncoupled. In addition, the heterochromatin defects in the *prp10*-1 mutant cannot rescued by expression of wild type RNAi genes, a*go1*^+^ and *hrr1*^+^, which contain introns. They further found an RNAi component, Cid12, interacts with splicing factors, including Prp10. They proposed that splicing factors interact with the RNAi machinery to directly regulate RNAi-mediated heterochromatin silencing (23). However, a comprehensive analysis of global splicing changes in these studies were missing, leaving open the possibility that splicing changes of a yet unknown heterochromatin factor might be responsible for the heterochromatin defects. Indeed, the study by Kallgren *et al*. has shown that splicing factor mutants, such as *cwf14*, result in the splicing defects in Ago1, which in turn lead to heterochromatin defects (25).

In this study, we performed the first comprehensive analysis on global splicing changes in a splicing mutant showing heterochromatin defects, which could potentially rule out any indirect influences. We show that deletion of *rbm10*^+^ has little effect on splicing, but leads to significant loss of silencing and H3K9 methylation in heterochromatin. In addition, Rbm10 interacts with a variety of heterochromatin factors, notably the HDAC Clr6 complex, and chromatin remodeling HIRA, FACT, CHD1 and Spt6 components. Furthermore, deletion of *rbm10*^+^ disrupts the association of the Clr6 complex with heterochromatin. These findings strongly suggest that Rbm10 is directly involved in heterochromatin formation. Rbm10-interacting splicing factor Cwf10 has been shown to be associated with pericentromeric transcripts (23). We propose that Rbm10 binds nascent heterochromatic transcripts, and serves as a platform to recruits the Clr6 complex and chromatin remodelers to heterochromatic region, which in turn promote heterochromatin assembly. Our study, together with previous findings, further suggests that different splicing subunits may play separate roles in regulation of heterochromatin formation.

### Role of Rbm10 in regulation of RNA splicing

In addition to its role in heterochromatin formation, our data suggest conserved function of Rbm10 in splicing. We detected that a plethora of Rbm10-interacting partners in fission yeast are from spliceosomal A and B complexes, which recapitulate the known interactions of human RBM10. Furthermore, overexpression of Rbm10 in fission yeast leads to massive intron retention, which can be seen as an equivalent to human RBM10’s function in promoting exon skipping in humans. These results suggest that fission yeast and human Rbm10 have retained similar function in RNA splicing.

However, we only observed the strong effect of Rbm10 on splicing in fission yeast when it is overexpressed. In the deletion mutant strain only very modest changes in splicing were detected, indicating a mild impact of Rbm10 on splicing at its endogenous expression level. We observed that Rbm10 expression level appears to be extremely low under normal cell growth condition. Given the fact that stoichiometric amounts of splicing factors are needed for splicing, our data indicate that Rbm10 does not serve as a general splicing factor under endogenous expression level. We speculate that higher expression of *rbm10*^+^ may be induced under specific environmental conditions, leading to changes of gene expression through altering RNA splicing patterns. Due to its strong toxicity in overexpression, it is likely that *rbm10*^+^ expression level is tightly controlled and its dual function might be regulated dose-dependently.

### Interactome of Rbm10

We show that Rbm10 interacts with around 850 proteins, with enrichment for splicing factors, HDAC, chromatin remodelers and transcription factors. These data strongly support the direct role of Rbm10 in heterochromatin formation and splicing. A recent study showed that a splicing factor in *Arabidopsis* also interacts with HDAC (57). Both of these processes are involved with various large protein complexes, explaining the high number of interacting partners. Especially the good correlation between fission yeast and human RBM10 in regard to its interaction with the spliceosome can be seen as a proof-of-principle experiment.

Rbm10 interacts with transcription factors, especially Byr3, which has the highest sequence coverage of the whole dataset. Since heterochromatin formation is also dependent on transcription during S phase, we speculate that this factor may play a role in mediating heterochromatin transcription. Its exact role in heterochromatin regulation still needs further characterization in future studies. Interestingly, we also found that Rbm10 interacts with multiple replication factors (Figure 5D). Replication factors have also been implicated in heterochromatin and centromere assembly in fission yeast (14,22,58–61).

### Comparison between fission yeast and human RBM10

The fact that Rbm10 interacts with the spliceosome and its overexpression results in intron retention indicates that the function of Rbm10 as a splicing factor is conserved between human and fission yeast. However, human RBM10 is expressed at substantially higher level than fission yeast Rbm10. This may be due to the fact that human RBM10 regulates exon skipping that generates alternative spliced proteins, but fission yeast Rbm10, on the other hand, is mainly responsible for intron retention, which can lead to premature block of transcription and RNA degradation via nonsense-mediated RNA decay (NMD) pathway (62). Overexepression and knock-down of human RBM10 lead to massive changes in gene expression (26,27). Overexepression of human RBM10 also results in nuclear condensation (63). However, it is still not clear whether the defects are evoked by secondary effects, or result from that fact that human RBM10 has a role in heterochromatin formation and gene silencing. Nevertheless, human RBM10 was found to be part of H2A deubiquitinases complex (36), which has been implicated in heterochromatin regulation (64). Future work is needed to elucidate whether human RBM10 has a dual function in the regulation of splicing and heterochromatin formation.

## Materials and Methods

### Fission yeast strains and genetic analysis

*rbm10*Δ, *rbm10-GFP* and *FLAG-HA-rbm10* strains were constructed via a PCR-based homologous recombination and verified by PCRs. *pREP1-FLAG-HA-rbm10* was constructed by inserting their respective cDNAs into NdeI/BamHI or SalI/SmaI sites in the plasmid. Genetic crosses were conducted following standard protocols (65). Fission yeast strains used in this study are listed in the Supplemental Table S1. For silencing assays, a series of 10-fold dilutions with a starting concentration of 2× 10^7^ cells/ml were spotted on the designated PMG (Pombe glutamate medium) media or the EMM (Edinburgh Minimal Medium) media with FOA and incubated at 30°C for 2-3 days.

### RNA Sequencing

RNA sequencing was performed as described (26). Briefly, poly (A) RNA was prepared from total RNA by two rounds of oligo (dT)_25_ Dynabeads (Invitrogen) purification. Following fragmentation at 94°C for 3.5 min, the RNA was converted to first strand cDNA using random hexmer primer and Supescript II (Invitrogen), followed by second strand cDNA synthesis with *Eschericcia coli* DNA pol I (Invitrogen) and RNAse H (Invitrogen). The barcoded sequencing library was prepared and sequenced on Illumina HiSeq for 1 × 100 cycles following the standard protocol.

### RNA-Seq data analysis

RNA-seq reads that pass the Illumina filter were aligned to the *S. pombe* genome reference sequences (version: ASM294v2) using TopHat with default mapping parameters (version 2.0.8) (66). The read count for each exon-exon and/or exon-intron junction as well as each gene was produced by custom Perl script. DESeq package was used to identify the differentially expressed genes between *rbm10*Δ cells and WT (67). Adjusted p.value < 0.05 was used as the threshold for determining significance. Fisher’s exact test was used to compare the splicing pattern of each exon (for exon skipping) or each intron (for intron retention) between *rbm10*Δ cells and WT. Adjusted p.value < 0.05 and |ΔPSI| (or |PIR|) > 10% was used as the threshold for determining significant splicing difference.

### ChIP

ChIP was performed by as previously described (68). Briefly, 50 ml of a log-phase yeast culture was cross-linked by adding 37% formaldehyde for 30 min. Cells were collected and sonicated by an Ultrasonic Processor. 1 μl of H3K9me antibody (Abcam ab1220) was used for immunoprecipitation. Immunoprecipitated DNA was purified using a PCR clean up column (Qiagen), and analyzed by PCR using primers listed in Supplementary Table S8. The experiments were replicated in two or three independent biological repeats.

### RT-PCR

RT-PCR was performed by as previously described with minor modifications (69). 500 ng starting material of total RNA was used per reaction. First strand synthesis was performed with Random Primer and Superscript II RT (Invitrogen) at 25°C for 10 min, then 42°C for 50 min and 70°C for 15 min. For the subsequent PCR, 1 μl from the reverse transcriptase reaction was used. Oligos used are listed in Supplementary Table S8.

### TAP-tag purification and mass spectrometry

Cells carrying *pREP1-FLAG-HA-rbm10* was induced for 24h in minimal media without thiamine. Cell were collected and lysed in lysis buffer with 4 μl/ml Benzonase by bead beating. Protein G beads coated with anti-FLAG-antibody were added to the cell lysate, and incubated for 1 hour at 4°C on rotation. After washing, the sample was eluted using 3 ×FLAG peptide (Sigma). The eluate was subsequently incubated with uMACS HA magnetic beads on ice for 30 min with occasionally mixing. The eluate with the beads was loaded on a uMACS column, and washed three times with wash buffer I and twice with wash buffer II. To prepare the sample for MS, proteins on the column were predigested by adding 25 μl 2M urea in 100 mM Tris-HCl pH 7.5, 1 mM DTT and 150 ng Trypsin for 30 min at RT. The sample was eluted by adding 2 times 50 μl 2M urea in 100 mM Tris-HCl, pH 7.5 and 5 mM iodocetamide. The sample was digested overnight at RT. 1 μl trifluoroacetic acid was added to stop the digestion. Mass spectrometry analysis was conducted in the Proteomics platform at Berlin Institute for Medical Systems Biology.

### Mass spectrometry Data processing and analysis

Raw data were analyzed using the MaxQuant proteomics pipeline (v1.4.0.5) and the built in the Andromeda search engine (Cox, Neuhauser et al. 2011) with the PomBase database, Carbamidomethylation of cysteines was chosen as fixed modification, oxidation of methionine and acetylation of N-terminus were chosen as variable modifications. The search engine peptide assignments were filtered at 1% FDR. The feature match between runs was left disabled and other parameters were left as default.

### Western blot analysis

Western blot was performed by as previously described (70). Briefly, cell extracts from exponentially growing cells were collected. Extracted proteins were separated on SDS-polyacrylamide gels and blotted onto PVDF membranes. Blots were probed with indicated antibodies.

### Co-immunoprecipitation (Co-IP)

Co-immunoprecipitation was conducted as described (61). Cells were collected and resuspended in 100 μl of 1 ×lysis buffer with protease inhibitors prior to lysis by bead beating. Lysates were incubated with lgG sepharose (GE Healthcare) at 4 °C for 2 hrs. After washing with lysis buffer three times, proteins were eluted in SDS loading buffer. Eluates were analyzed by Western blotting using a commercial anti-GFP antibody (Abcam, ab290).

### Microscopy

Cells were imaged using the DeltaVision System (Applied Precision, Issaquah, WA). Images were taken as z-stacks of 0.2-μm increments with an oil immersion objective (×100).

Standard DAPI staining and analysis methods for fission yeast nuclei were used.

## Data access

All genomic data were available at the GEO database under the accession number GSE93906.

## Supplementary Information

Supplementary Data are available Online.

## Author contributions

F.L. and W.C. designed experiments and wrote the manuscript; M.W. and H.B. performed experiments with assistance from Q.G., and H.H.. G.M. and S.K. performed the mass spectrometry analysis.

## Acknowledgments

We thank Qianhua Dong and Jinpu Yang for critical reading of the manuscript.

## Ethics Declarations

### Ethics approval and consent to participate

Not applied.

### Consent for publication

Not applied.

